# Effect of *Thiobacillus* and superabsorbent on essential oil components in *Thyme* species

**DOI:** 10.1101/430983

**Authors:** Pouneh Pouramini, Mohammad Hossein Fotokian, Hossein Dehghan, Goetz Hensel

## Abstract

Optimal nutrition along with non-stress conditions has a significant impact on the quantity and quality of essential oil in medicinal plants. The objective of this research was to examine the possibility of improving the quantity and quality of essential oil in *thyme* through nutrition of the seedlings using *Thiobacillus* bio-fertilizer and treatment by Superabsorbent. For this purpose, seedlings of two *thyme* species (*Thymus vulgaris* and *Thymus daenensis*) were sown in pots and exposed to different levels of *Thiobacillus* and superabsorbent. Results of Gas Chromatography Mass Spectrometry (GC/MS) revealed that the main compounds detected for *T. vulgaris* were thymol (31.5%), *p*-cymene (23.4%), *γ*-terpinene (13.9%), linalool (38.3%) and carvacrol (2.7%) while the main compounds of *T. daenensis* were thymol (51.2%), *o*-cymene (12.9%), *γ*-terpinene (4.5%), linalool (1.7%) and borneol (3.1%). Furthermore, the application of *Thiobacillus* had a significant effect on *α*-pinene content (p = 0.05). Moreover, the interaction between superabsorbent and *Thiobacillus* significantly changed the percentage of thymol, borneol, and caryophyllene. In conclusion, the mean of essence components in *T. vulgaris* was more than *T. daenensis* with the exception of thymol and caryophyllene.

## 1. Introduction

The genus *Thymus*, with about 215 species, is one of the eight most important genera of the Lamiacea family. *Thyme*, a plant native to the Mediterranean region (Spain, Italy, France, Greece, etc.), has long been used as a source of essential oil and other constituents (e.g. thymol, flavanoid, caffeic acid and labiatic acid) derived from the different parts of the plant. In addition to their numerous traditional uses, the plant (herb) and its essential oil have found diverse applications in pharmacy and medicine [1]. *Thymus vulgaris* is a perennial medicinal plant, cultivated worldwide for a lot of uses like culinary, cosmetic and medical purposes. This species has special activities such as antispasmodic, expectorant, antiseptic, antimicrobial and antioxidant [2,3]. *Thymus daenensis* is native to Iran [4].

Depending on the growing conditions (different environments, climates, soil, geographical location) *Thyme* can provide various therapeutic properties [5]. Plant growth, quality, and quantity of active ingredients mainly depends on genetic manipulation while environmental factors play a major role by leading to changes in the growth, quality and quantity of active substances such as alkaloids, glycosides, essential oils [6].

*Thyme* contains 0.8 to 2.6% (generally 1%) of essential oils which is primarily comprised (20 to 80%) of *phenols, monoterpenes* (e.g. *p-cymene* and *y-terpinen*), and *alcohols* (e.g. *linalool, a-terpinene* and is *thujan-4-ol*). In general, *thymol* constitutes the highest content of phenolic compounds while *carvacrol* is a minor part of the essential oil of *Thyme* [7]. In a study by Barazandeh and Bagherzadeh [8] investigating the essential oil of the aerial parts of *Thymus daenensis*, GS/MS results showed that among the 43 indicated components, *thymol* (73.9%), *carvacrol* (6.7%), *para-cymene* (4.6%), *β-bisabolene* (1.5%) and *terpinene 4-L* (1.4%) were the main constituents.

Bio-fertilizers are of great importance and play the largest role in increasing the productivity and maintaining soil fertility in sustainable agriculture [9]. Bio-fertilizers are microbial inoculants consisting of living cells of microorganism like bacteria, algae and fungi alone or combination which can help in increasing crop productivity. Organic manures can serve as alternative to mineral fertilizers for improving soil structure, soil nutrients, soil physical and chemical characteristics, increase soil fertility and plant plants resistance to diseases and salt stress, decrease plant diseases, improving crop growth, development, yield and quality through directly synthesizing hormones, antibodies and secondary metabolites [10,11]. through organic and biological methods plays a key role in increasing qualitative and quantitative herbal and essential oil yield [12].

In many regions of Iran, soil type is calcareous with high pH. Therefore, some nutrients such as phosphorus, iron and zinc are stabilized and non-absorbable for plants. In such situations, *Thiobacillus* bacteria can reduce the pH of the soil in the presence of sulfur resulting in the solubility of nutrients around the root area [13].

These previously mentioned issues have increased along with limited water resources in the arid and semi-arid ecosystems of Iran [14]. Water stress has many adverse effects on the plant’s processes such as reduction in the leaf area index, the number of leaves and shoots dry weight [15,16]. Medicinal plants are not accepted and water stress reduces the quantity and quality of active ingredients while these herbs need to perfect vegetative and reproductive growth in order to combine the ingredients [16]. Superabsorbents are able to reduce the need to be irrigated by up to 33% with no side effects on the soil and the environment [14]. Superabsorbent is natural or synthetic contain over 99% water. Hydrogels have been defined as polymeric materials which exhibit the ability of swelling in water and retaining a significant fraction (>20%) of water within their structure, without dissolving in water [17]. Superabsorbent hydrogels are increase soil capacity to hold water, nutrients and improve the plant’s growth and yield [18]. Although the application of superabsorbent and *Thiobacillus* have been separately studied, to our knowledge, their combined effects have not yet been examined on medicinal plants. Therefore, this research was conducted to investigate the possibility of improving the quantity and quality of essential oil in *Thyme* through nutrition of the seedlings by *Thiobacillus* bio-fertilizer and increase of available water through the application of a superabsorbent.

## 2. Materials and methods

### 2.1 Experimental condition

The experiment was conducted in a greenhouse on a research farm of an Agricultural Engineering Park affiliated with the Ministry of Agriculture in Jihad, Karaj, Iran (56.5 ° East longitude and 35.47 ° N latitude) during 2012. Plant growth was carried out in a greenhouse at 24/18°C day/night mean temperature and 65% air relative humidity. Properties of used soil are presented in Table 1.

**Table 1.** Selected physical and chemical properties of soil used in this study.

| Tissue | P<br>[ppm] | K<br>[ppm] | Total<br>Nitrogen<br>[%] | Organic<br>matter<br>[%] | Calcium<br>carbonate<br>equivalent<br>[%] | pH | EC<br>[ds/m] | Percent<br>saturation<br>[%] |
| --- | --- | --- | --- | --- | --- | --- | --- | --- |
| Loamy | 38 | 344.7 | 0.22 | 2.19 | 7.5 | 8.26 | 1.32 | 45 |

### 2.2 Treatments

A factorial experiment using a completely randomized design with two replications was performed. The experiment consists of three factors of *Thiobacillus* (two levels of present and absent), superabsorbent (three levels of 0, 0.5 and 1 g k^-1^ soil) and *Thyme* (*Thymus vulgaris* and *Thymus daenensis*).

### 2.3 Herb processing and essential oil isolation

The seeds were planted on 5 April 2011 and germinated after 17 days. The first flower was visible on 13 July, while the plants were harvested at a full flowering phase on 30 August, air-dried at 25°C in the shade and subsequently well crushed. The essential oil was extracted by hydro distillation for 120 min using a Clevenger-type system [19]. Collected essential oils were stored in dark flasks and kept at –18°C for GC/MS analysis.

### 2.4 Statistical analysis

Statistical analyses were conducted using SPSS version 16.0. A normality test of collected data residuals was conducted. Analysis of variance (ANOVA) was performed followed by Duncan’s multiple range test to compare the effects of main and/or sub-main factor levels combination. Pearson’s correlation coefficients were used to detect associations between dependent measures. Statistical significance was concluded with a *P*-value less than 0.05.

## 3. Results and discussion

### 3.1 *Comparison of Thymus vulgaris* and *Thymus daenensis* essential oil composition

The *Thymus vulgaris* essential oil included 31 constituents which represented 96.7% of the total analyzed oil. The main compounds included thymol (31.5%), *p*-cymene (23.4%), *γ*-terpinene (13.9%), linalool (3.4%), thymol methyl ether (3.6%) and carvacrol (2.7%). Further compounds and their constituents are presented in Table 2. In contrast, the *Thymus daenensis* species contained 33 compounds in the essential oil which represented 96.19% of the total analyzed oil. The main compounds included thymol (51.18%), O-cymene (12.87%), γ- Terpinene (4.48%), linalool (1.68%) and borneol (3.09%). The major differences are in the thymol content as well as monoterpenes (Table 2, grey highlighted). While some compounds are present only in one species, some other compounds show only minor differences between both species.

**Table 2.** Compounds identified in *Thymus vulgaris and Thymus daenensis*

| No. | Compound | *Thymus vulgaris* |  | *Thymus daenensis* |  | Class |
| --- | --- | --- | --- | --- | --- | --- |
|  |  | RT [min] | [%] | RT [min] | [%] |  |
| 1 | α-Thujene | 11:12 | **1.51** | 11:05 | **0.90** | MH |
| 2 | α-Pinene | 11:44 | **1.17** | 11:35 | **0.68** | MH |
| 3 | Camphene | 12:10 | **0.78** | 12:05 | **0.52** | MH |
| 4 | β-Pinene | 13:49 | **0.36** | 13:44 | **0.30** | MH |
| 5 | Octen-3-ol | - | - | 13:86 | **0.23** | Others |
| 6 | β-Myrcene | 14:39 | **3.32** | 14:32 | **0.88** | MH |
| 7 | Phellandrene |  |  | 14:91 | **0.18** | MH |
| 8 | α-Phellandrene | 15:23 | **0.44** | - | - | MH |
| 9 | *p*-Cymene | 16:90 | **23.40** | - | - | MH |
| 10 | α-Terpinene | - | - | 15:60 | **1.14** | MH |
| 11 | *o*-Cymene | - | - | 16:31 | **12.87** | MH |
| 12 | 1,8-Cineole | - | - | 16:52 | **1.94** | MO |
| 13 | γ-Terpinene | 18:63 | **13.94** | 17:95 | **4.48** | MH |
| 14 | β-Terpineol | 18:80 | **0.87** | 18:31 | **0.84** | MO |
| 15 | α-Terpinolene | 19:50 | **0.25** | 19:24 | **0.15** | MH |
| 16 | Linalool | 20:47 | **3.38** | 20:06 | **1.68** | MO |
| 17 | Borneol | 23:34 | **1.02** | 23:73 | **3.09** | MO |
| 18 | Terpinen-4-ol | 23:85 | **0.84** | - | - | MO |
| 19 | Borneol | 24:49 | **1.61** | - | - | MO |
| 20 | α-Terpineol | - | - | 25:76 | **3.30** | MO |
| 21 | Dihydro carvone | - | - | 26:16 | **0.13** | MO |
| 22 | Thymol methyl ether | 26:72 | **3.58** | 26:47 | **1.19** | MO |
| 23 | Carvacrol methyl ether | 27:10 | **1.65** | 26:89 | **0.58** | MO |
| 24 | Thymoquinone | - | - | 28:59 | **3.02** | MO |
| 25 | Thymol | 30:62 | **31.50** | 30:64 | **51.18** | MO |
| 26 | Carvacrol | 31:13 | **2.66** | - | - | MO |
| 27 | Caryophyllene | 35:06 | **1.55** | 34:96 | **2.41** | SH |
| 28 | Aromadenderene | - | - | 35:67 | **0.17** | SH |
| 29 | α-Humulene | - | - | 36:28 | **0.28** | SH |
| 30 | Geranyl propionate | - | - | 37:00 | **0.12** | Others |
| 31 | Geranyl propionate | 37:13 | **0.46** | - | - | Others |
| 32 | α-Amorphene | 37:26 | **0.12** | - | - | SH |
| 33 | Germacrene D | 37:46 | **0.13** | - | - | SH |
| 34 | Viridiflorene | - | - | 37:95 | **0.15** | SH |
| 35 | Cubebol | 37:99 | **0.10** | - | - | SH |
| **36** | *β*-Bisabolene | - | - | 38:47 | **0.42** | SH |
| **37** | *γ*-Cadinene | 38:77 | **0.18** | 39:05 | **0.16** | SH |
| **38** | *δ*-cadinene | 39:13 | **0.43** | - | - | SO |
| **39** | Thymohydro quinone | - | - | 40:82 | **1.18** | SO |
| **40** | Caryophelen oxide | 41:54 | **0.75** | 41:52 | **2.13** | SO |
| **41** | Eudesmol | 42:87 | **0.09** | - | - | SO |
| **42** | *α*-epi-Cadinol | 43:68 | **0.21** | - | - | SO |
| **43** | *α*-Cadinol | 44:17 | **0.10** | - | - | SO |
| **44** | (E)-14-Hydroxy-9-epi-caryophyllene | 44:81 | **0.13** | - | - | SO |
| **45** | *α*-epi-Bisabolol | - | - | 45:20 | **0.12** | Others |
| **46** | Hexadecanoic acid | 54:87 | **0.13** | 54:84 | **0.13** | Others |
| **47** | Abiatatriene | - | - | 60:20 | **3.22** | MH |
| **Total Identified** |  |  | **96.67** |  | **96.19** |  |

MH – monoterpene hydrocarbons; MO – oxygenated monoterpenes; SH – sesquiterpene hydrocarbons; SO – oxygenated sesquiterpenes

The components identified in *Thymus vulgaris* are in accordance with a study analysing material from Saudi Arabian markets [20]. Since *Thymus daenensis* had a higher thymol content compared to *Thymus vulgaris* previous studies focused on this species. Our results find thymol still the main component of the *Thymus* oil but the quantities are below previous published data. The reason for that might be the different soil characteristics because the studies have been performed in different parts of Iran [4,8].

Extractable oil composition strongly depends on the method used. In a study by Charles and Simon [21] comparison of different extraction methods showed that hydro distillation on basil lead to the greatest number of constituents with the highest number of volatile constituents in the organic solvent extract. Therefore, hydro distillation appears to be an excellent method for extracting the essential oil of basil as it results in good yield, good recovery of essential oil constituents, is less labor-intensive, and is simpler and faster than steam distillation.

### 3.2 Treatment with superabsorbent and Thiobacillus on essential oils content

Analysis of variance of some components of thyme essential oil is presented in Table 3. The difference between two thyme species was significant (*P* ≤ 0.01) for all of the quantities of the compounds (Table 4) while the superabsorbent effect was not significant for any compound. The *Thiobacillus* effect was only significant for *α*-Cymene (*P* ≤ 0.05). In the case of treatment interactions, it was found that the superabsorbent × *Thiobacillus* interaction had a significant effect on borneol and caryophylene compounds. Furthermore, the species × *Thiobacillus* interaction was significant for thymol. The triple interaction among species × superabsorbent × *Thiobacillus* had a significant effect on borneol and caryophylene.

**Table 3.** Analysis of variance of the effects of superabsorbent and *Thiobacillus* on the essential oil of two species of thyme

| S.O.V | df | Mean squares |  |  |  |  |  |  |  |  |  |
| --- | --- | --- | --- | --- | --- | --- | --- | --- | --- | --- | --- |
| | | α-Thujene | α-Pinene | Camphene | β-Myrcene | *p*-Cymene | γ-Terpinene | Linalool | Borneol | Thymol | Caryophyllene |
| S | 1 | 0.562<sup>**</sup> | 1.321<sup>**</sup> | 0.702<sup>**</sup> | 11.323<sup>**</sup> | 2110.034<sup>**</sup> | 441.438<sup>**</sup> | 19.706<sup>**</sup> | 3.683<sup>**</sup> | 2545.542<sup>**</sup> | 28.423<sup>**</sup> |
| SA | 2 | 0.024<sup>ns</sup> | 0.058<sup>ns</sup> | 0.038<sup>ns</sup> | 0.061<sup>ns</sup> | 7.561<sup>ns</sup> | 11.952<sup>ns</sup> | 0.005<sup>ns</sup> | 0.232<sup>ns</sup> | 74.919<sup>ns</sup> | 0.251<sup>ns</sup> |
| T | 1 | 0.022<sup>ns</sup> | 0.003<sup>ns</sup> | 0.001<sup>ns</sup> | 0.018<sup>ns</sup> | 40.755<sup>*</sup> | 0.832<sup>ns</sup> | 0.438<sup>ns</sup> | 0.061<sup>ns</sup> | 1.322<sup>ns</sup> | 0.295<sup>ns</sup> |
| T×SA | 2 | 0.026<sup>ns</sup> | 0.054<sup>ns</sup> | 0.034<sup>ns</sup> | 0.001<sup>ns</sup> | 10.054<sup>ns</sup> | 4.751<sup>ns</sup> | 0.238<sup>ns</sup> | 0.481<sup>*</sup> | 11.094<sup>ns</sup> | 0.559<sup>*</sup> |
| S×SA | 2 | 0.021<sup>ns</sup> | 0.030<sup>ns</sup> | 0.010<sup>ns</sup> | 0.075<sup>ns</sup> | 0.319<sup>ns</sup> | 11.882<sup>ns</sup> | 0.338<sup>ns</sup> | 0.001<sup>ns</sup> | 41.655<sup>ns</sup> | 0.263<sup>ns</sup> |
| S×T | 1 | 0.055<sup>ns</sup> | 0.107<sup>ns</sup> | 0.018<sup>ns</sup> | 0.002<sup>ns</sup> | 25.058<sup>ns</sup> | 0.695<sup>ns</sup> | 0.096<sup>ns</sup> | 0.034<sup>ns</sup> | 433.359<sup>*</sup> | 0.112<sup>ns</sup> |
| S×T×SA | 2 | 0.009<sup>ns</sup> | 0.026<sup>ns</sup> | 0.036<sup>ns</sup> | 0.008<sup>ns</sup> | 22.564<sup>ns</sup> | 5.611<sup>ns</sup> | 0.183<sup>ns</sup> | 0.384<sup>*</sup> | 61.169<sup>ns</sup> | 0.469<sup>*</sup> |
| Error | 11 | 0.013 | 0.027 | 0.028 | 0.022 | 8.132 | 3.251 | 0.317 | 0.089 | 43.097 | 0.108 |
| CV % | - | 26.4 | 25.1 | 29.2 | 17.5 | 16.9 | 29.2 | 29.4 | 24.6 | 11.6 | 16 |

Symbols: S, Species; SA, Superabsorbent; T, *Thiobacillus*.
\*, \*\* significant differences at $P \leq 0.05$ and $P \leq 0.01$ , respectively.

**Table 4.** Mean comparisons between two species for essence components by Duncan’s multiple range test

| Species | α-Thujene | α-Pinene | Camphene | β-Myrcene | *p*-Cymene | γ-Terpinene | Linalool | Borneol | Thymol | Caryophyllene |
| --- | --- | --- | --- | --- | --- | --- | --- | --- | --- | --- |
| *T. vulgaris* | 0.583 | 0.887 | 0.743 | 1.529 | 26.181 | 9.192 | 2.823 | 1.603 | 46.297 | 0.964 |
| *T. daenensis* | 0.277 | 0.418 | 0.401 | 0.156 | 7.428 | 0.615 | 1.011 | 0.8197 | 66.895 | 3.141 |

-Refer to variance analysis (Table 3), the differences between two *Thymus* species is statistically significant at $P \leq 0.01$ .

A review of the literature regarding the application of *Thiobacillus* showed that there was little information about the use on medicinal plants. In a recent study, the effect of Bio-fertilizer with superabsorbent on the yield and essential oil content of lemon verbena (*Lippia citriodora* Kunth.) were analysed. The authors could show that sole application of treatments was useful, but their combined use was more effective to increase dry mass and oil yield of lemon verbena [18]. In another study in dragonhead, the effect of chemical fertilizer and microbial inoculum application on the yield and nutrients content of *Dracocephalum moldavica* L. seeds was evaluated. In a field experiment, the highest yield was obtained by chemical fertilizers, while seed inoculation with sulfur oxidizing bacteria (*Thiobacillus strains*) improves the iron nutrient content of the seeds [21]. Also, in the presence of sulfur, *Thiobacillus* increased nutrient uptake, growth and production of *Melissa officinalis* oil compared to the control [12].

Razban and Pirzad [16] investigated the effect of superabsorbent on *Matricaria chamomilla* growth under different irrigation regimes. They reported that superabsorbent was able to compensate for the reduction of biomass yield due to lack of water.

### 3.3 Relations between the main compounds of essential oil

Pearson’s correlation coefficients between traits showed a significant positive association between the numbers of traits (*P* ≤ 0.01) (Table 5) suggesting these may be due to pleiotropic effects of genes controlling the same pathways, close linkage, and/or epistasis effects.

**Table 5.**
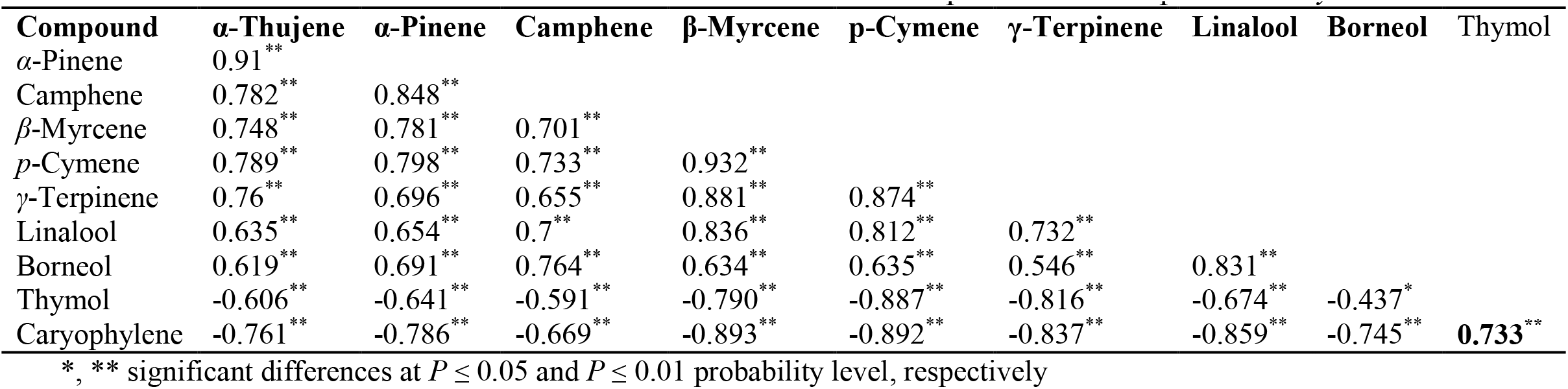
Pearson’s correlation coefficients between some essential components of two species of *Thyme*

| Compound | α-Thujene | α-Pinene | Camphene | β-Myrcene | *p*-Cymene | γ-Terpinene | Linalool | Borneol | Thymol |
| --- | --- | --- | --- | --- | --- | --- | --- | --- | --- |
| α-Pinene | 0.91<sup>**</sup> | | | | | | | | |
| Camphene | 0.782<sup>**</sup> | 0.848<sup>**</sup> |  |  |  |  |  |  |  |
| β-Myrcene | 0.748<sup>**</sup> | 0.781<sup>**</sup> | 0.701<sup>**</sup> | | | | | | |
| *p*-Cymene | 0.789<sup>**</sup> | 0.798<sup>**</sup> | 0.733<sup>**</sup> | 0.932<sup>**</sup> |  |  |  |  |  |
| γ-Terpinene | 0.76<sup>**</sup> | 0.696<sup>**</sup> | 0.655<sup>**</sup> | 0.881<sup>**</sup> | 0.874<sup>**</sup> | | | | |
| Linalool | 0.635<sup>**</sup> | 0.654<sup>**</sup> | 0.7<sup>**</sup> | 0.836<sup>**</sup> | 0.812<sup>**</sup> | 0.732<sup>**</sup> |  |  |  |
| Borneol | 0.619<sup>**</sup> | 0.691<sup>**</sup> | 0.764<sup>**</sup> | 0.634<sup>**</sup> | 0.635<sup>**</sup> | 0.546<sup>**</sup> | 0.831<sup>**</sup> |  |  |
| Thymol | -0.606<sup>**</sup> | -0.641<sup>**</sup> | -0.591<sup>**</sup> | -0.790<sup>**</sup> | -0.887<sup>**</sup> | -0.816<sup>**</sup> | -0.674<sup>**</sup> | -0.437<sup>*</sup> |  |
| Caryophyllene | -0.761<sup>**</sup> | -0.786<sup>**</sup> | -0.669<sup>**</sup> | -0.893<sup>**</sup> | -0.892<sup>**</sup> | -0.837<sup>**</sup> | -0.859<sup>**</sup> | -0.745<sup>**</sup> | **0.733<sup>\*\*</sup>** |

\*, \*\* significant differences at $P \leq 0.05$ and $P \leq 0.01$ probability level, respectively

### 3.4 Interaction between factors

#### 3.4.1 Superabsorbent × Thiobacillus interaction

Combined levels of the superabsorbent × *Thiobacillus* interaction were significant for borneol and caryophyllene (Table 4 and Figure 1; *P* ≤ 0.05). The maximum amount of borneol was observed in S_0_T_0_ treatment, although there was no significant difference in the amount of borneol in the treatment S_1_T_0.5_. Although the maximum amount of caryophyllene was observed in S_0.5_T_0.5_, the difference of this treatment with S_0_T_0.5_ and S_1_T_0_ treatments was not statistically significant (Figure 1). The differences among superabsorbent levels were not statistically significant for essential components. One reason for this may be the low difference among levels of superabsorbent. With the increase of the content in superabsorbent treatment levels, the differences among these levels may be significant for essence components.

**Figure 1.**
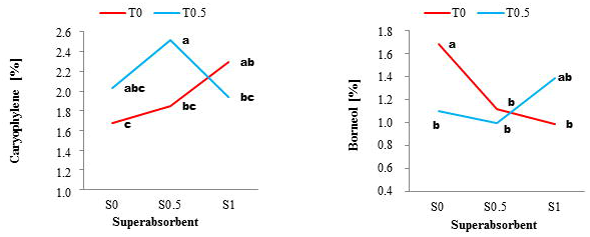
Caryophyllene and borneol change trend across superabsorbent and *Thiobacillus* interaction levels over the *Thymus* species. Values with different letters in each graph are significantly different according to Duncan’s Multiple Range Test (P ≤ 0.05).

#### 3.4.2 Species × Thiobacillus interaction

The interaction between species × *Thiobacillus* was significant for thymol (*P* ≤ 0.05; Table 4). Using *Thiobacillus* in *Thymus daenensis* led to a significant reduction in the amount of thymol and an increase in the amount of thymol in *Thymus vulgaris* (Figure 2).

**Figure 2.**
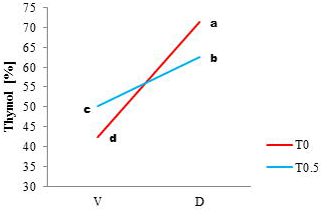
Mean comparisons of Thymol under the interaction of species × *Thiobacillus*. V and D present *Thymus vulgaris* and *Thymus daenensis*, respectively. Values with different letters are significantly different according to Duncan’s Multiple Range Test (P ≤ 0.05).

#### 3.4.3 The triple interaction among species × superabsorbent ×Thiobacillus

Significant differences among combinations of this interaction were observed only for Borneol and caryophyllene (*P* ≤ 0.05) (Table 6). The maximum amount of caryophyllene was obtained with *Thymus daenensis* in the S_0.5_T_0.5_ treatment, with no statistically significant differences of this treatment with S_1_T_0_. Application of superabsorbent and *Thiobacillus* did not improve the amount of borneol. The amount of borneol in *Thymus vulgaris* was significantly higher than *Thymus daenensis*.

**Table 6.** Comparison interaction of species, superabsorbent and *Thiobacillus* on borneol and caryophylene on *Thymus vulgaris* and *Thymus daenensis*

| Interactions | Caryophyllene [%] | Borneol [%] |
| --- | --- | --- |
| V × S0 × T0 | 0.9934<sup>d</sup> | 2.289<sup>a</sup> |
| V × S0 × T0.5 | 0.9528<sup>d</sup> | 1.301<sup>bcd</sup> |
| V × S0.5 × T0 | 0.8876<sup>d</sup> | 1.281<sup>bcd</sup> |
| V × S0.5 × T0.5 | 1.124<sup>d</sup> | 1.621<sup>abc</sup> |
| V × S1 × T0 | 0.8859<sup>d</sup> | 1.278<sup>bcd</sup> |
| V × S1 × T0.5 | 0.9458<sup>d</sup> | 1.849<sup>ab</sup> |
| D × S0 × T0 | 2.351<sup>c</sup> | 1.081<sup>cde</sup> |
| D × S0 × T0.5 | 3.112<sup>bc</sup> | 0.9007<sup>cde</sup> |
| D × S0.5 × T0 | 2.82<sup>c</sup> | 0.9435<sup>cde</sup> |
| D × S0.5 × T0.5 | 3.92<sup>a</sup> | 0.3738<sup>e</sup> |
| D × S1 × T0 | 3.715<sup>ab</sup> | 0.6984<sup>de</sup> |
| D × S1 × T0.5 | 2.93<sup>c</sup> | 0.9215<sup>cde</sup> |

-Treatment symbols: T<sub>0</sub>, 0.0 g *Thiobacillus*; T<sub>0.5</sub>, 0.5 g *Thiobacillus*; S<sub>0</sub>, 0.0 g superabsorbent; S<sub>0.5</sub>, 0.5 g superabsorbent; S<sub>1</sub>, 1.0 g superabsorbent; V, *Thymus vulgaris*; D, *Thymus daenensis*.
-Values with different letters in a column are significantly different according to Duncan's Multiple Range Test (P ≤ 0.05)

The maximum amount of borneol was observed in the S_0_T_0_ treatment, although there was no significant difference in the amounts found in the S_1_T_0.5_ treatment. Although the maximum amount of caryophyllene was observed in S_0.5_T_0.5_, but the difference of this treatment with S_0_T_0.5_ and S_1_T_0_ treatments was not statistically significant (Figure 1).

The interaction between species × superabsorbent was not significant for all traits, while for species main factor it was significant at *P* ≤ 0.01. Referring to an acceptable amount of coefficient of variation (CV %), it is concluded that non-significance of some parts of variation for essence components sources is not due to the amount of experimental error.

The correlation between thymol with others was negatively significant, with the exception of a positive correlation with caryophylene. In conclusion, thymol is the most important part of essence component in *Thymus*; any trying to increase thymol content will decrease other essence components.

## 4. Conclusion

The interaction between *Thiobacillus* and superabsorbent on oil and essential oil yield was significantly greater for borneol and caryophyllene. Furthermore, *Thymus vulgaris* oil yield was higher than *Thymus daenensis*. Despite the fact that *Thymus vulgaris* is not native to Iran, its high value in the pharmaceutical, cosmetic, health and culture, food and export will encourage its cultivation. In addition, the variance analysis of our data showed the amount of all components in *T. vulgaris* are much higher than *T. daenensis*, with the exception of thymol and caryophyllene. Hence, in the cultivation of *Thymus* for the production of essence components, cropping of *T. vulgaris* is recommended.

